# nanoCLAMP potently neutralizes SARS-CoV-2 and protects K18-hACE2 mice from infection

**DOI:** 10.1101/2023.04.03.535401

**Authors:** Quentin Pagneux, Nathalie Garnier, Manon Fabregue, Sarah Sharkaoui, Sophie Mazzoli, Ilka Engelmann, Rabah Boukherroub, Mary Strecker, Eric Cruz, Peter Ducos, Ana Zarubica, Richard Suderman, Sabine Szunerits

**Affiliations:** Univ. Lille, CNRS, Centrale Lille, Univ. Polytechnique Hauts-de-France, UMR 8520 - IEMN, F-59000 Lille, France; Univ Lille, CHU Lille, Laboratoire de Virologie ULR3610, F-59000 Lille, France; Centre d’Immunophénomique, Aix Marseille Université, Inserm, CNRS, PHENOMIN, Marseille, France; Pathogenesis and Control of Chronic and Emerging Infections, INSERM, EFS, Univ Antilles, Univ Montpellier, Laboratoire de Virologie, CHU Montpellier, France; Department of Biochemistry, University of Wisconsin-Madison, Madison, Wisconsin, 53706, USA; Nectagen, Inc., 2002 W 39th Ave, Kansas City, KS 66103, USA; Regis University, Colorado, USA; Celerion, Inc., 621 Rose Street, Lincoln, NE 68502, USA

**Author notes:** Corresponing authors.

**Keywords:** nano-CLostridial Antibody Mimetic Proteins (nanoCLAMP) spike protein, K18-hACE2 transgenic mice, SARS-CoV-2 infection

## Abstract

Intranasal treatments, combined with vaccination, has the potential to slow mutational evolution of virusues by reducing transmission and replication. Here we illustrate the development of a SARS-CoV-2 receptor binding domain (RBD) nanoCLAMP and demonstrate its potential as an intranasally administered therapeutic. A multi-epitope nanoCLAMP was made by fusing a pM affinity single-domain nanoCLAMP (P2710) to alternate epitope binding nanoCLAMP, P2609. The resulting multimerised nanoCLAMP P2712 had sub-pM affinity for the Wuhan and South African (B.1.351) RBD (K_D_ < 1 pM), and decreasing affinity for the Delta (B.1.617.2) and Omicron (B.1.1.529) variants (86 pM and 19.7 nM, respectively). P2712 potently inhibited ACE2:RBD interaction, suggesting its utility as a therapeutic. With an IC_50_ = 0.4 ± 0.1 nM obtained from neutralization experiments using pseudoviral particles as well as patient cultured SARS-CoV-2 samples, nanoCLAMP P2712 protected K18-hACE2 mice from SARS-CoV-2 infection, reduced viral loads in the lungs and brains, and reduced associated upregulation of inflammatory cytokines and chemokines. Together, our findings warrant further investigation into the development of nanoCLAMPs as effective intranasally delivered COVID19 therapeutics.

## 1. Introduction

While vaccines remain the most important weapon in the fight against the COVID-19 pandemic, there is an unmet need for the development of intranasally administered pre- and post-infection prophylactic treatments to prevent infection (Hadjichrysanthou et al, 2022, Ratcliffe et al, 2021). Along with vaccination, therapeutic monoclonal antibodies were one of the initial envisioned strategies in treating SARS-CoV-2 infection. Four intramuscularly delivered anti-SARS-CoV-2 mAb products (bamlanivimab plus etesevimab, casirivimab plus imdevimab, sotrovimab, and bebtelovimab) received Emergency Use Authorizations for the treatment of outpatients with mild to moderate COVID-19 and were based on the reduction of viral load (Cameroni et al, 2022, Chi et al, 2020, Kuramochi et al, 2022, Weinrich et al, 2020). These systemic neutralizing antibodies mainly target the Receptor-Binding Domain (RBD) of the trimeric Spike protein (Renn et al, 2020) that decorates the surface of the coronavirus and plays a pivotal role during viral entry by first binding to Ace2 displayed on epithelial cells 15(Dai et al, 2020). Since the main port of entry of SARS-CoV-2 into the body is the ciliated epithelium lining the nose with one of the highest concentration of Ace2 receptors of any tissue in the body, the deployment of countermeasures to that area has the potential to block the initial attachment of the virus to Ace2 receptors, pre-empting infection. Importantly, this strategy does not allow the virus to replicate and mutate in the host, as it does even in many vaccinated individuals whose immune systems effectively eliminate the virus. Since intramuscular injection of the current vaccines induces mainly IgG protection in the lower respiratory tract, there is a new interest in nasal or mucosal vaccinations to promote IgA antiviral activity at the point of entry and hypothesized to reduce replication comparatively (Ratcliffe et al, 2022). In combination with a nasal vaccine that produces IgA protection in the nasal passage, nasal treatments that block the initial attachment of the virus to host cells could drastically reduce transmission, replication, and by extension, mutation.

Several non-protein based nasal treatments for COVID-19 are under development and include povidone-iodine, nitric oxide, ethyl lauroyl arginate hydrochloride, and astodrimer sodium (Ratcliffe et al. 2021). Protein-based biologics employed as nasal treatments include an engineered IgM with prophylactic efficacy when delivered intranasally in rodents (Ku et al, 2021). Monoclonal antibodies are the most widespread affinity protein therapeutics due to their high specificity, affinity, and safety, with RBD targeting neutralizing monoclonal antibodies in development since the start of the pandemic (Cameroni et al. 2022, Imai et al, 2023, Lin et al, 2023, Renn et al. 2020, Wang et al, 2023). The use of therapeutic anti-spike monoclonal IgG, while effective, is challenging due to possible dramatic change of activity of anti-SARS-CoV-2 mAbs against specific variants and subvariant. Encouragingly, however, a broadly neutralizing antibody (35B5) was shown recently to neutralize all known variants of concern when administered intranasally (Lin et al. 2023). While monoclonal antibodies remain successful protein therapeutics, they are expensive to produce and are prone to degradation and aggregation in extreme conditions (Sécher et al, 2022). Further, their development requires immunization of an animal, adding valuable time to their development. The cost for a single dose of intranasal mAb, which could possibly require several milligrams of antibody, is likely to be prohibitive for mass distribution, especially in developing countries. Therefore, low cost antibody mimetics are emerging as an attractive alternative as nasal prophylactics.

Nanobodies, and other antibody mimetics are being developed as alternatives to traditional antibodies for nasal prophylactics (Case et al, 2021, Chonira et al, 2023, Li et al, 2022, Pymm et al, 2021, Titong et al, 2022, Wu et al, 2021, Wu et al, 2022). These single domain binding proteins can be produced cheaply in microbial hosts, have better engineering and stability profiles, and have the potential to address the high cost associated with mAb nasal prophylactics. A purely synthetic, potent miniprotein using structural design alone to develop picomolar binders to the spike RBD was designed by Case et al (Case et al. 2021). The well characterized ankyrin repeat domain-based Darpins are also being developed as intranasal prophylactics (Chonira et al. 2023). Wu et al (Wu et al. 2021) report a potent bispecific nanobody that protects mice against SARS-CoV-2 infection via intranasal administration. The nanobody NIH-CoVnb-112 was effectively nebulized and demonstrated effective reductions viral burden, and lung pathology in a Syrian hamster model of COVID-19. More recently Wu et al (Wu et al. 2022) demonstrated short term instantaneous prophylaxis and treatment in hAce2 mice infected with the delta variant of SARS-CoV-2 2 with nanobody NB-22-Fc. Encouragingly, a single dose of intranasal Nb22 protected mice even when administered 7 days prior to infection with the Delta variant.

While nanobodies dominate this field, their development typically originates in immunized Camelidae, followed by cloning and development of the binding domain (the nanobody), adding time to their development. Furthermore, many camelid-derived nanobodies contain disulfide bonds, making them susceptible to reducing environments, forcing them to be produced in the periplasm of *E. coli*, or in secretion expression systems, and precluding their engineering with sulfhydryl reactive probes and reagents. Recently a new class of antibody mimetics called nanoCLAMPs (nano-CLostridial Antibody Mimetic Proteins) has been described (Suderman et al, 2017), with attractive properties for the development of an intranasal treatment. These 15 kDa protein binders can be screened from a synthetic, naïve phage display library of 1 × 10^10^ variants for high specificity, low nM affinity binders to targets in 6 weeks, and produced and purified cheaply from the cytosol of *E. coli* with yields above 200 g L^-1^. They are naturally devoid of cysteines, are easily refolded following chemical denaturation with 6 M GuHCl, 0.1 N NaOH, or DMF when conjugated to solid support, and have high thermal stability (T_m_ > 65 C), highlighting their stability in harsh environments. Importantly, they are easily multimerized, making them well suited countermeasures to quickly mutating targets, since escape mutants must simultaneously mutate two epitopes.

Here we describe the development of two nanoCLAMPs with low nanomolar affinity for two distinct epitopes on the Wuhan RBD, the affinity maturation of one of them, and finally their fusion into a multi-epitope-binding, sub-picomolar affinity nanoCLAMP, P2712. We characterized this protein’s ability to inhibit human Ace2:RBD binding *in vitro* and further investigated this activity by demonstrating the neutralization potential of P2712 using pseudovirus bearing the SARS-CoV-2 spike trimer. The neutralizing potential of P2712 on COVID-19 patient samples was also tested. Finally, we demonstrate for the first time the efficacy of a nanoCLAMP as a potential intranasally delivered therapeutics using a SARS-CoV-2 susceptible K18-hACE2 mouse model. Reduced viral load in lungs and brain and reduced upregulation of inflammatory cytokines and chemokines underline the potential of nanoCLAMPs as effective intranasally delivered COVID19 therapeutics.

## 2. Results

### 2.1. High-Affinity SARS-CoV-2 RBD-Binding Single-Domain Antibody Mimetic nanoCLAMP P2710

With the aim to obtain nanoCLAMPs that potently neutralize SARS-CoV-2, the synthetic nanoCLAMP phage display library NL-21 (1×10^10^ variants) was screened for binders to the SARS-CoV-2 Wuhan strain (Wuhan-Hu-1) spike protein receptor binding domain (RBD). After 3 rounds of phage panning, we screened 96 random clones by semELISA and identified 41 unique nanoCLAMPs specific to recombinant RBD and ranked them according to off-rate using biolayer interferometry (BLI) with recombinant RBD (data not shown). Clones with poor monodispersity were carefully excluded, as soluble aggregates artificially increased the apparent affinity due to increased avidity. For this reason, we purified the monodisperse monomer of each nanoCLAMP by size exclusion chromatography, and tested the monodispersity prior to kinetic and functional analyses (*SI Appendix, Fig. S4*). The nanoCLAMP P2632 was determined to be the lead candidate with a *K*_D_ = 3.3 nM for recombinant Wuhan RBD determined by Bio-layer interferometry (BLI) on a streptavidin sensor chip to which biotinylated-Wuhan RBD was immobilized (*SI Appendix, Fig. S2*). NanoCLAMP P2632 was affinity matured by constructing and screening a saturation mutagenesis library for three additional rounds of phage panning, resulting in nanoCLAMP P2710, with *K*_D_ ≤ 1 pM for the Wuhan-RBD (**Figure 1**). The low *K*_D_ is notably due to the nondetectable dissociation rate, *k*_off_, by BLI, where single digit pM and fM affinities cannot be reliably determined. NanoCLAMP P2710 has one of the highest affinities for RBD of the reported single-domain antibody mimetics (without multimerizing), with comparable affinity to some of the nanobodies from immunized Llamas reported by Güttler et al. (Güttler et al, 2021). While the affinity of nanoCLAMP P2710 to the U.K. variant (B.1.1.7) remains comparable to that of the Wuhan-RBD, affinities towards the S. African RBD (B.1.351), Delta RBD (B.1.617.2) and Omicorn-RBD (B1.1.529) are markedly lower, indicating that P2710 affinity was affected by mutations in the RBD.

**Figure 1.**
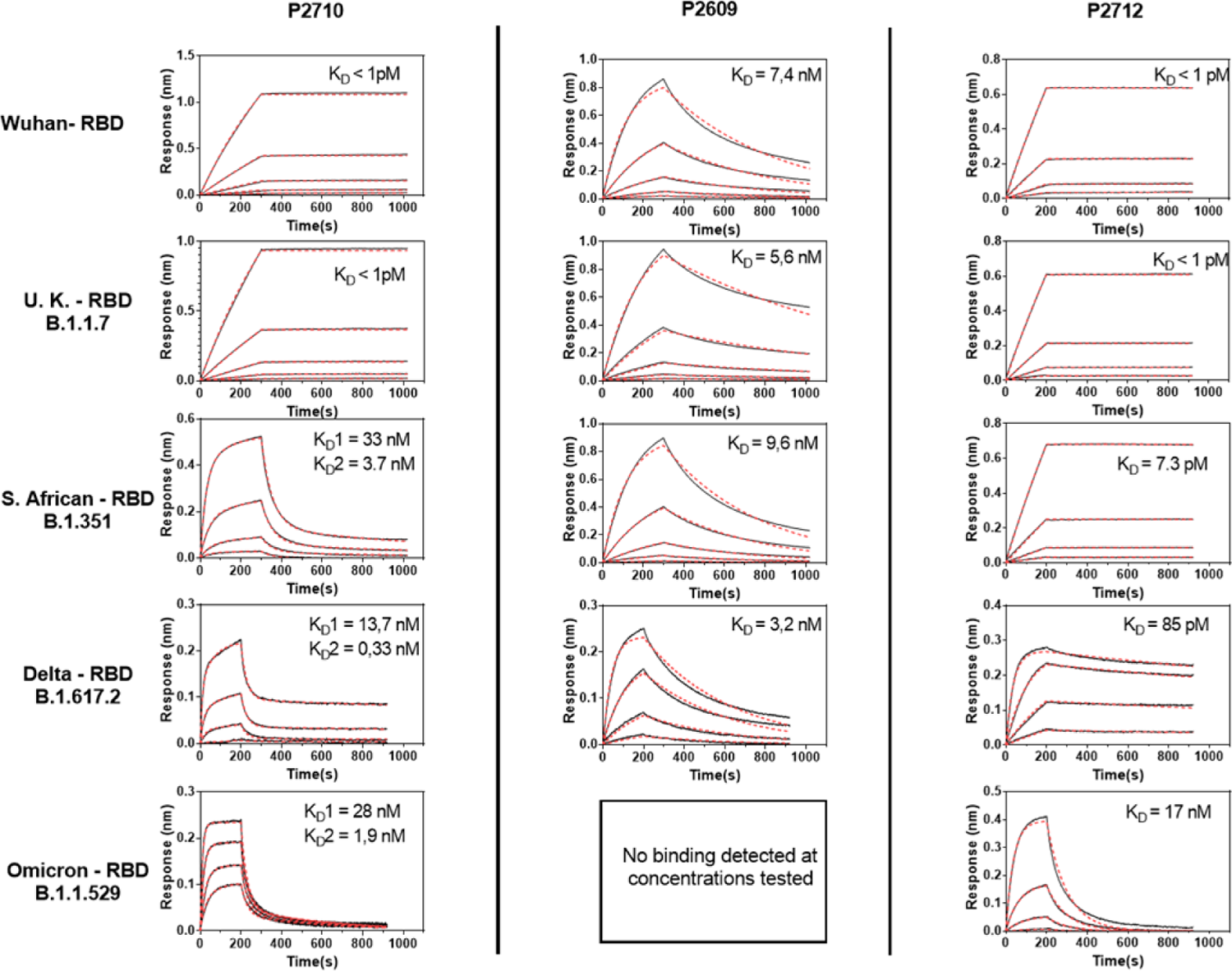
Binding affinities of nanoCLAMPs to SARS-CoV-2 RBD from VOCs. BLI sensorgrams showing binding of nanoCLAMPs P2710, P2609, and P2712 (fusion of P2609 and P2710) to recombinant SARS CoV 2 RBDs from different variants of concern. Blck lines depict binding data and red lines depict 1:1 binding model fit, except in cases of P2710 vs Beta-RBD, Delta-RBD and Omicron-RBD, which was modeled with heterogenous ligand fit. Analyte concentrations were 10, 3.33, 1.11, 0.37 nM for all except: P2712 vs Omicron-RBD, P2710 vs Delta-RBD and P2609 vs Delta-RBD where 30,10,3.33,1.11 nM were used, and P2710 vs Omicron-RBD with 60,30,15,7.5 nM.

### 2.2. High-Affinity Multimeric RBD-Binding Single-Domain Antibody Mimetic nanoCLAMP P2712

Escape mutants pose a challenge for therapeutic approaches dependent on inhibiting RBD:Ace2 interaction. Given the prevalence of circulating RBD mutations and their impact on affinity, we hypothesized that fusing two or more nanoCLAMPs targeting different epitopes on the RBD would result in a higher affinity binder, and reduce the effects of single RBD mutations on affinity. The affinity of nanoCLAMP P2609 for Wuhan-RBD was in the nanomolar (*K*_D_= 7.4 nM) range (**Figure 1**), but had similar affinity and binding kinetics for U.K., S. African and Delta RBD (popular variants), suggesting this binder contacted regions not perturbed by mutations. Since the omicron variant had not yet emerged, we reasoned that the P2609 epitope might be conserved among variants and decided to determine whether it could be combined with P2710 to enhance binding to all variants.

Epitope binning revealed that P2609 binds a different epitope than P2710, and both nanoCLAMPs could bind RBD at the same time (**Figure 2A**). Attachment of biotinylated nanoCLAMP P2710 to a streptavidin modified BLI surface shows strong binding to Wuhan-RBD as expected. Addition of nanoCLAMP P2710 resulted in no additional binding, while added binding was observed with the addition of nanoCLAMP P2609. Based on these data, P2710 was fused with P2609 via a 13 residue GS linker, resulting in nanoCLAMP P2712 (6His-P2710-linker-P2609). P2712 expressed at high levels as expected and was purified to homogeneity (*SI Appendix, Figs S3)*. The thermal stability of the individual nanoCLAMPs and the P2712 fusion was tested using differential scanning fluorimetry (*SI Appendix, Fig. S4*). The thermal stability of P2710 and P2609 (Tm = 72 and 76 °C, respectively) was only slightly diminished in the fusion, with a Tm = 67 °C. We then measured the binding affinity of P2712 for RBD variants by BLI, and found that fusing P2710 and P2609 displayed low-pM binding for Wuhan, UK, and South African variants, and a KD of 85 pM and 17 nM for the Delta and Omicron variants, respectively (**Figure 1**). Since P2609 had constant affinity for the Wuhan, UK, and S. African variants, we anticipated it would retain its affinity for the Delta and Omicron variants. Interestingly, while it retained similar affinity for the Delta variant, its binding was nearly abolished by the omicron variant (**Figure 1**). We were surprised to see that P2710 retained some binding to the highly mutated omicron variant, imparting at least nM affinity to the P2712 fusion for the Omicron variant.

**Figure 2.**
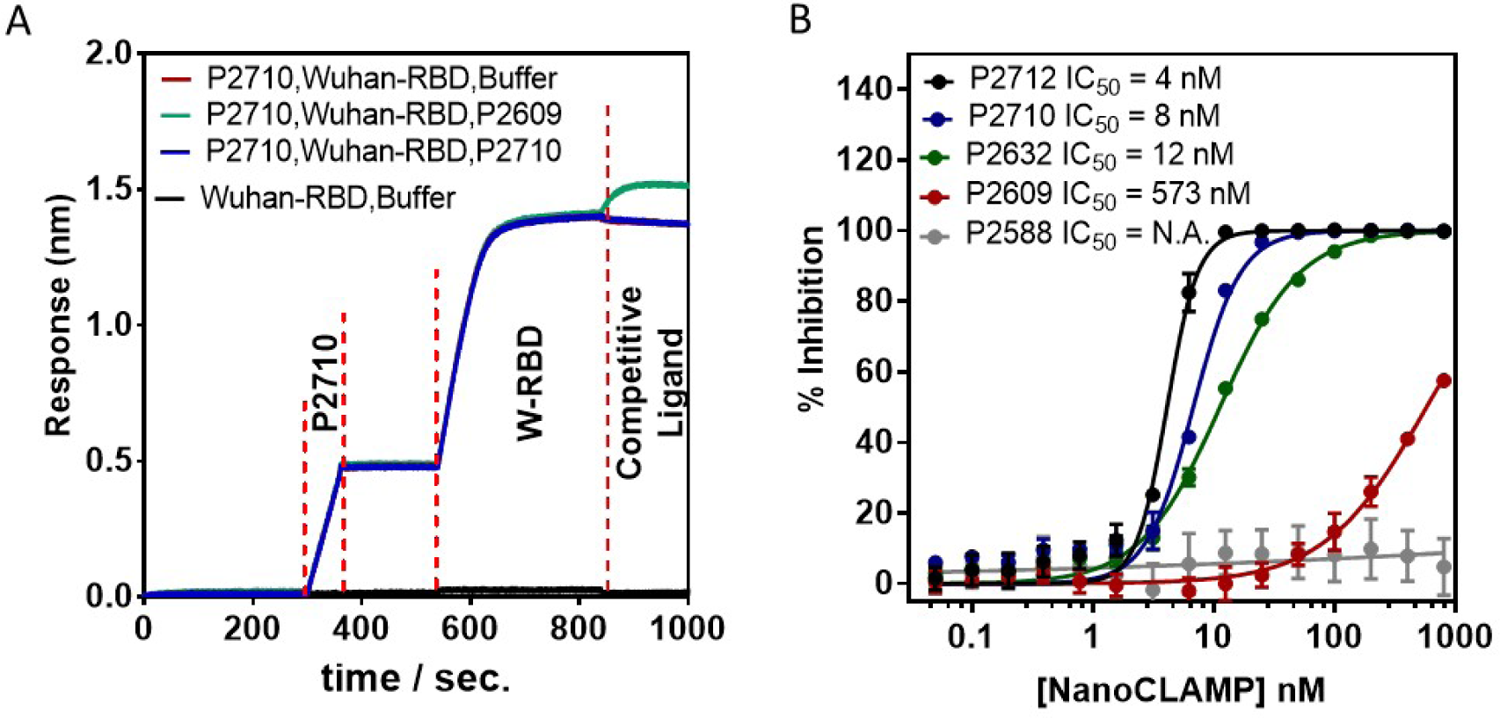
NanoCLAMP P2712 interactions and inhibitions: (A) Multimeric NanoCLAMP P2712 derived of P2710 and P2609 targeting different RBD epitopes: BLI sensogram of a streptavidin coated sensor showing loading of sensor with biotinylated P2710 (red), baseline establishment, binding of Wuhan RBD, and competitive binding of P2609 (green), P2710 (blue, dotted), or buffer (black). (B) Protein-A coated wells were coated with Wuhan RBD-Fc and incubated with biotinylated ACE2 that had been incubated with serial dilutions of nanoCLAMPs. Remaining ACE2 was detected with Streptavidin-HRP and developed with TMB-ELISA substrate. Data are represented as mean +/- SD, with n=3 biological replicates. IC50 values were calculated using Softmax Pro nonlinear regression 4 parameter fit.

As the pandemic has highlighted not only the need for a broad biological toolbox for therapeutics, but also for diagnostic tools, we tested whether P2712 could function as an affinity ligand in surface plasmon resonance (SPR), as well as the demonstrated BLI (above). SPR experiments demonstrated that biotinylated nanoCLAMP P2712 can indeed serve as a bioreceptor recognizing the different RBD variants (*SI Appendix, Fig. S5*). Kinetic analysis of immobilized, biotinylated P2712 binding to the Wuhan, U.K. S. African, Delta and Omicron variant RBDs yielded K_D_ in close agreement with the BLI data (**Figure 1**).

Although the mutational drift of SARS-CoV-2 was causing a decline in the affinity of P2712 for the RBD, we decided to continue to test whether nanoCLAMPs might be good candidates for intranasal therapeutics or prophylactics, reasoning that demonstration of the proof of principle could enable future therapeutic developments with different nanoCLAMPs, perhaps to more conserved regions of the Spike protein. With this aim in mind, we began testing the neutralization potential of P2712.

### 2.3. Inhibition of ACE-2:RBD interaction by nanoCLAMP P2712

Intranasal delivery of prophylactics and therapeutics against respiratory viruses is an attractive option because it delivers the agent directly to the viral point of entry and potentially blocks the virus from binding to and entering epithelial cells. It also has the potential to limit systemic exposure which could drastically reduce off-target effects. We hypothesized that the small size and high thermal, proteolytic, and chemical stability of nanoCLAMPs might make them good candidates for intranasally formulated therapeutics and prophylactics. As a first step to evaluate the therapeutic potential of nanoCLAMP P2712 and take advantage of its high affinity for the RBD, we tested in a competitive ELISA whether P2712 could inhibit the binding of the Wuhan-RBD domain of the SARS-CoV-2 spike protein to the extracellular domain of recombinant human ACE-2 receptor (**Figure 2B**). A non-RBD-binding nanoCLAMP, P2588, was included as a negative control and showed no inhibition of the interaction at any concentration tested, as expected. Although P2632 and P2609 both have similar affinity for Wuhan-RBD, it is clear that P2632, and it’s affinity-matured derivative P2710, inhibits the RBD:ACE2 interaction more potently than P2609, with IC_50_ values of 12 and 8 nM, respectively. The fact that P2609 did not inhibit RBD:ACE2 binding suggested that this binder might interact outside the region affected by the mutations in the SARS-CoV-2 variants of concern. P2712, the fusion of P2710 and P2609, was clearly the most potent inhibitor of the interaction, with an IC_50_ of 4 nM, and was further investigated for its neutralization potential.

### 2.4 Cytotoxicity of nanoCLAMP 2712 to Vero E6 cells and cell internalization studies

The cell toxicity of nanoCLAMP P2712 was established after 24 h incubation on Vero E6 cells, one of the most widely used cell lines for the proliferation and isolation of severe acute respiratory syndrome coronaviruses, as they contain an abundance of ACE2 receptors (Puray-Chavez et al, 2021). This receptor is expressed in lung epithelial cells as well as endothelial cells lining the nasal tract, arteries, veins, capillaries, small intestine, testes, renal tissue and cardiovascular tissue and is required for SARS-CoV-2 virus entry into the host cell. Infection of the host cell also relies on priming of the SARS-CoV-2 spike protein by the transmembrane serine protease (TMPRSS) (Hoffmann et al, 2020). The cytotoxicity was evaluated using cell viability assessment by the resazurin assay, based on the conversion of nonfluorescent dye to a fluorescent molecule by mitochondrial and cytoplasmatic enzymes. nanoCLAMP P2712 is nontoxic to Vero E6 cells up to high concentration (400 μg/mL) when incubated for 24 h (**Figure 3A**). To understand eventual uptake mechanism, Vero E6 cells were fixed and stained with P2712-ATTO488. The endocytosis percentage was quantitatively evaluated using flow cytometry by treating Vero E6 cells with 100 μg mL^−1^ of P2712 for 4 h at 37 °C. As 96 % of fluorescence remained in the initial R1 fluorescence population (**Figure 3B**) no internalized in Vero E6 cells of nanoCLAMPs occurs to a larger extend as could be expected.

**Figure 3.**
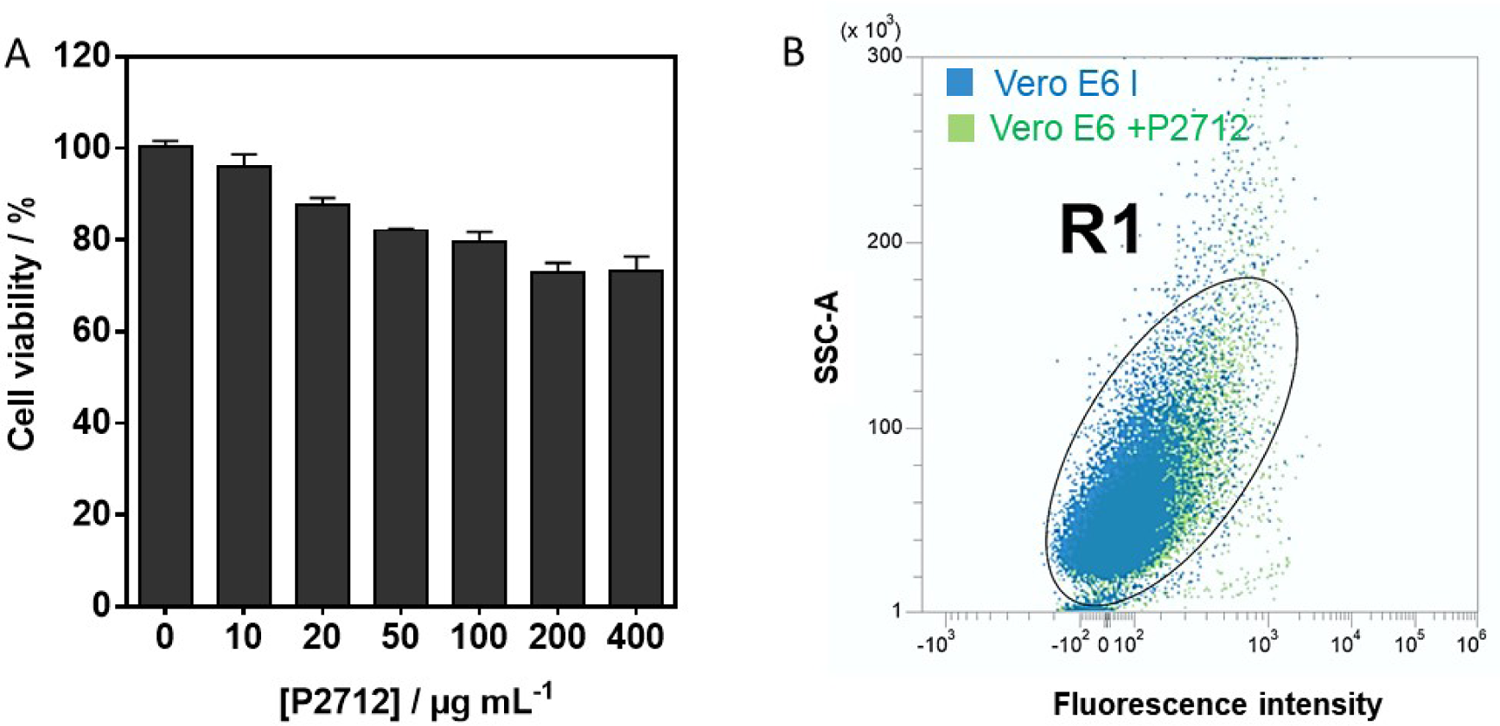
nanoCLAMP 2712 effect on Vero E6 cells: (A) Viability of Vero E6 cells treated with P2712 at different concentrations: Vero E6 cells were grown in 96-well plates (15 × 10^3^ cells/well) with 100 μL of culture medium containing increasing concentration of P2712 for 72 h. The results, expressed as percentage of viability, are the mean value of three independent experiments with each treatment performed in triplicate. Negative control: without P2712. (B) Fluorescence analysis with flow cytometry of Vero E6 cells untreated (Blue) and treated with 25.6 nM of P2712-ATTO488 for 4 h at 37 °C (Green).

### 2.5 nanoCLAMP P2712 neutralizes both SARS-CoV-2 pseudovirus and patient derived SARS-CoV-2 isolates

To test whether nanoCLAMP P2712 would recognize their targets in the context of the entire envelope Spike trimer, we tested their performance in a pseudovirus neutralization assay utilizing lentivirus pseudo typed with Wuhan SARS-CoV-2 Spike trimer (**Figure 4A**). Not unexpectedly, P2609 did not completely neutralize the pseudovirus in the concentration range tested. Full dose-response neutralization curves were generated for P2632, P2710, and P2712, with increasing neutralization IC_50_ of 5.4, 1.3, and 0.33 nM, respectively. Encouraged by these results, we tested P2712’s ability to neutralize other pseudovirus variants (**Figure 4B**). The the U.K. variant was neutralized with an IC_50_ of 0.21 nM in line with the Wuhan variant. Surprisingly and in contrast to the affinity results on RBD (**Figure 1**) P2712 did not significantly neutralize the S. African pseudoviral particles.

**Figure 4.**
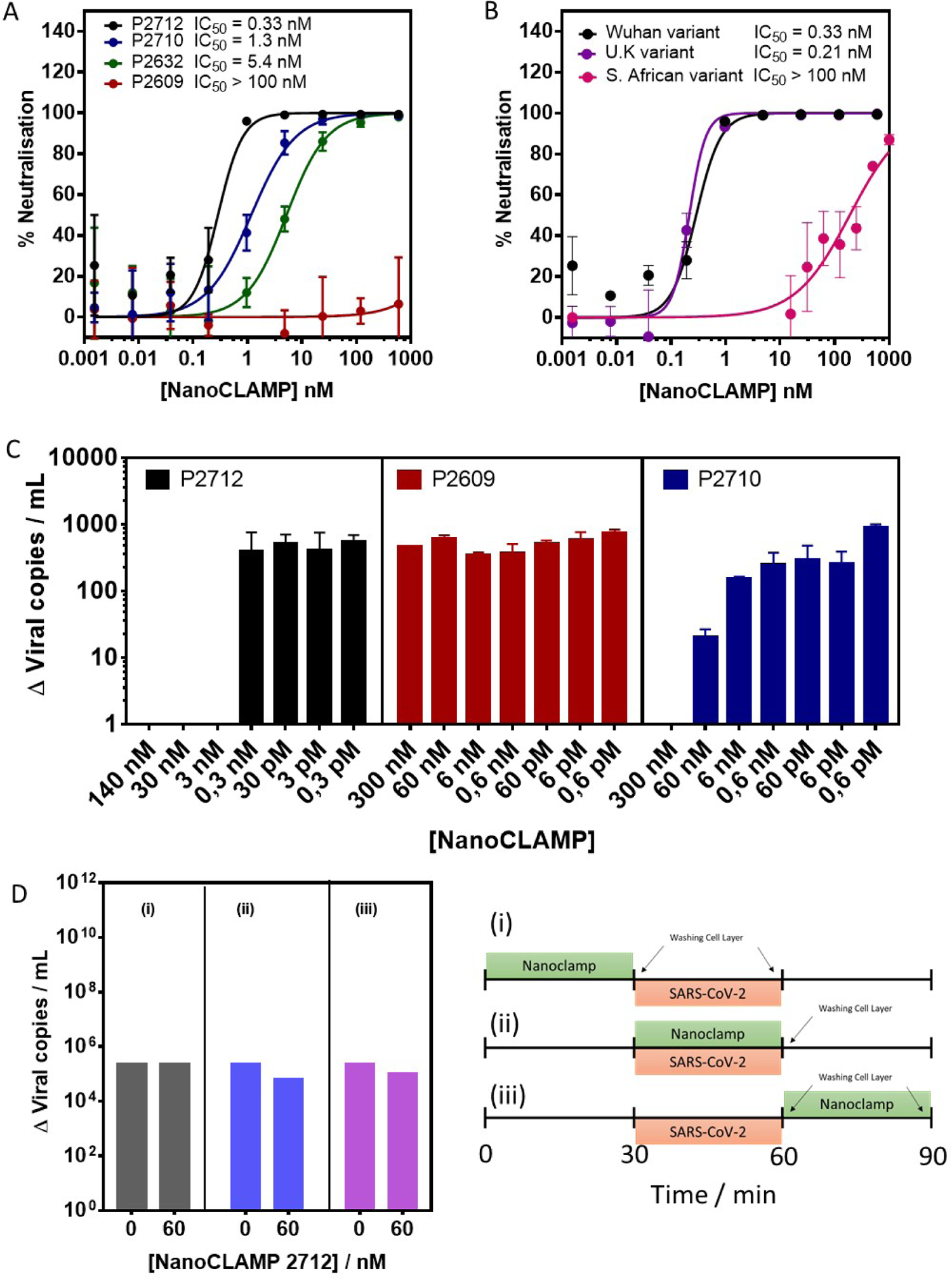
Neutralization and inhibition potency of nanoCLAMPs. (A) Neutralization of SARS-CoV-2 pseudotyped lentivirus infection of ACE-2 expressing HEK293T cells with RBD-specific nanoCLAMPs. Neutralization of Wuhan-Hu-1 PSV with nanoCLAMPs P2632 and P2609, which bind separate epitopes; the affinity matured version of P2632 (P2710) and the dual-epitope-binding fusion of P2710 and P2609 (P2712). (B). Neutralization of U.K. (B.1.1.7) and S. African (B.1.351) PSV by P2712 in comparison to Wuhan variant (same as Fig. 4a,black line). (C) Viral Inhibition assay using different nanoCLAMPs. Viral RNA in the culture supernatants were quantified by RT-qPCR after 5 days of incubation of Vero E6 cells with different concentrations of nanoCLAMPs and SARS-CoV-2 viral particles at 7.3 ×10^7^ viral particles mL^-1^ (incolum). (D) Mechanistic investigations: nanoCLAMP P2712 at 60 nM was added at different time points for 30 min during infection of VeroE6 cells with SARS-CoV-2, as shown in the graphs.

To assess the potential of nanoCLAMP P2712 to prevent viral entry, a SARS-CoV-2 neutralization assay was set up in parallel (**Figure 4C**). A patient-derived SARS-CoV-2 Wuhan isolate was incubated with varying concentrations of P2712 and then applied to monolayers of Vero 6 cells. Infectivity and neutralization were quantified five days post-infection by measuring viral RNA by RT-qPCR The initial SARS-CoV-2 concentration was 7.3×10^7^ viral particles mL^-1^.While no neuralization effects were observed for nanoCLAMPs 2609 (**Figure 4C**, nanoCLAMP P2712 reached the inoculum level at 3 nM concentration with nanoCLAMP 2710 showing neutralization onset at higher concentrations of 300 nM in line with results in **Figure 4A**. However using SARS-CoV-2 Delta and Omicron isolates, no neutralization effects were observed in the tested concentration range (data not shown).

We further investigated the mechanism of action of P2712 on SARS-CoV-2 Wuhan isolate infection by performing a time-on-addition assay. P2712 was added at different time points during infection (**Figure 4D**). Strong inhibition of infection was observed only when P2712 was mixed with the virus before being incubated with the Vero E6 cells suggesting that the entry step of the virus is blocked.

### 2.6 Intranasal administration of P2712 is highly efficacious as a therapeutic treatment of K18-hACE2 transgenic mice infected with Wuhan-Hu-1 SARS-CoV-2

Based on the success of the studies on pseudovirus and patients samples, we next assessed the efficacy of nanoCLAMP P2712 on an authentic SARS-CoV-2 challenge model using K18-hACE2 transgenic mice. Ten mice per group (8-to-10-week-old female) were administered a single, daily 30 µL intranasal dose of 2.25 mg/kg P2712 or an equimolar dose of negative control nanoCLAMP P2570 (corresponding to 1.12 mg/kg with a 30µL does) for 3 days, 6h before infection to day 2 after infection (**Figure 5A**). The mice were intranasally challenged with 2.5 × 10^4^ plaque-forming units (PFU) of SARS-CoV-2 per mouse, and change of body weight changes (**Figures 5B**), survival (**Figure 5C**) and clinical score (**Figure 5D**) monitored until the study end point at day 14. At day 5, 4 mice were sacrificed for tissue analysis.

**Figure 5.**
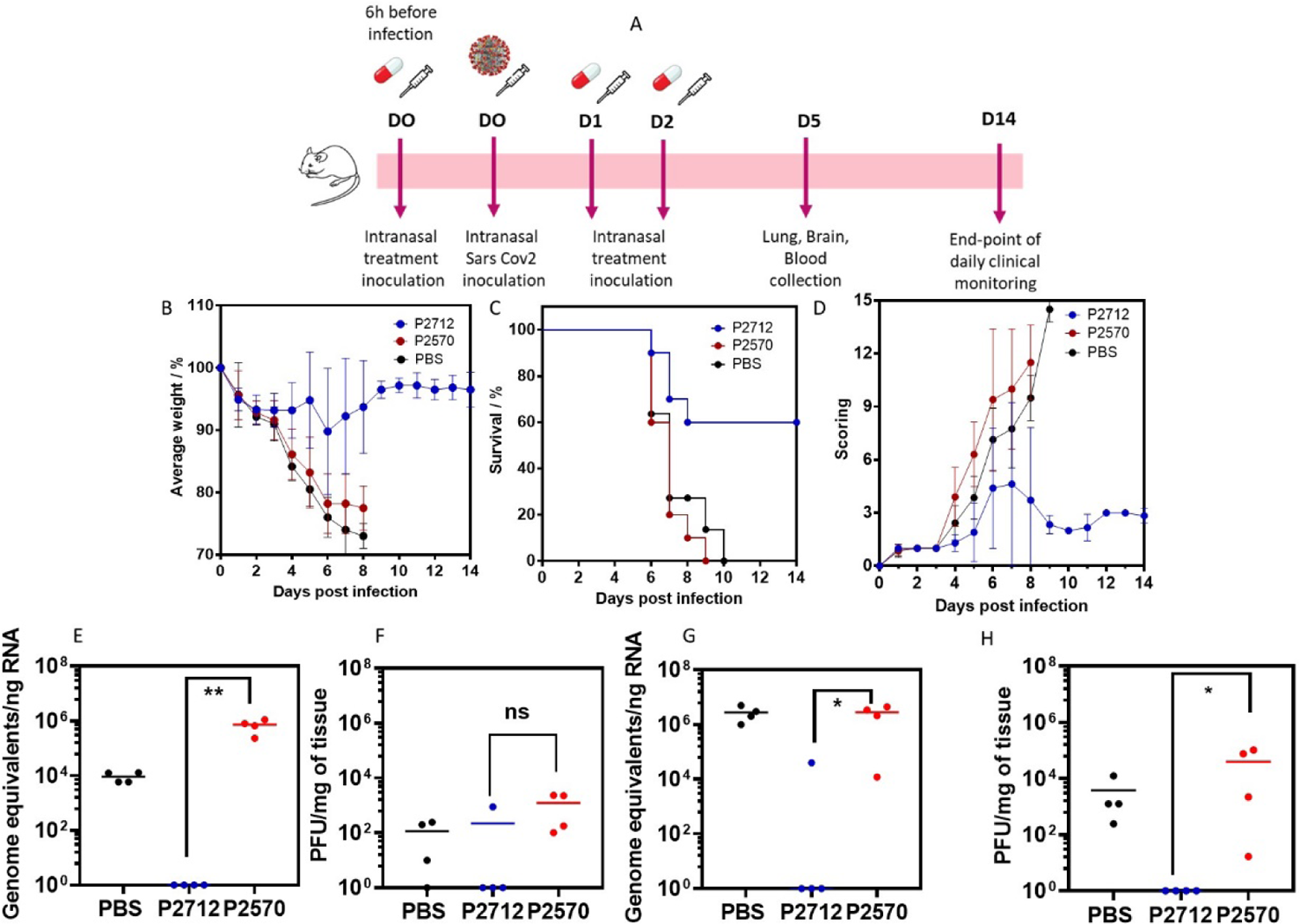
SARS-CoV-2 infection in nanoCLAMP treated K18-hACE2 mice with viral burden and virtal titer. (A) 8-to-10-week-old female K18-hACE2 mice were infected with SARS-CoV-2 and treated by PBS, mock or therapeutic compound nanoCLAMP P2570 and P2712 respectively as indicated on depicted schema; (B) Weight change as a function of time and represented as mean per therapeutic group (C) Survival follow-up (Mantel-Cox test) was daily evaluated until 14 days post infection; (D) Clinical score is monitoring and represented daily until end point of experiment at 14 days post infection as mean per therapeutic group (means from 10 individuals for each group treated with 5mg/kg nanoCLAMP). (E-H) SARS-CoV-2 viral burden and titer (qRT-PCR, TCID50) in infected and treated K18-hACE2 animals by PBS, mock (P2570) or therapeutic (P2712) compound at 5-day post infection. (E) Viral RNA level detected by RT-qPCR in lungs. (F) viral charge detected by TCID50 in lungs of SARS-CoV-2-infected K18-hACE2. (G) Viral RNA level detected by RT-qPCR in brain. (H) Viral charge detected by TCID50 in brains of SARSCoV-2-infected K18-hACE2 mice. (Each plotted dot represents the viral burden and titer at day 5 post-infection for an individual animal and bars represent median and standard deviation. Data analysis has been done by GraphPad, Mann-Whitney test.

At day 8 post infection (dpi), (before any mice had died); mice treated with the negative control, nanoCLAMP P2570, lost over 20% of their initial body weight and showed increasing clinical score (conjunctivitis, lethargy, hunched appearance, respiratory difficulties). All these mice succumbed to infection or were humanely euthanized due to exceeding humane endpoints previously defined. In contrast, the nanoCLAMP P2712 group showed reduced body weight loss (around 5-10%) and limited clinical score compared to nanoCLAMP P2570. Only 40% of the mice in P2712 group succumbed to infection or reached humane endpoint and were humanely euthanized. These mice showed over 20% body weight loss and high global clinical score. The remaining 60% showed no major changes in body weight and low grade of clinical score suggesting protection effect of P2712 on associated pathology.

After viral challenge with an infectious dose of 2.5×10^4^ PFU, the levels of viral RNA present in lungs and brains was assessed by RT-qPCR at day 5 post infection. Viral load was significantly decreased in lung and brain homogenates of mice treated with P2712 antibody (**Figures 5E,G**). In parallel, viral charge in lungs and brains was assessed by TCID50 measurements (**Figures 5F,H**). Differences were observed in viral charge between both groups, going from 100 to 10^3^ PFU/mg of tissue in lung, and from 100 to 10^5^ PFU in brain. The viral load in K18 hACE2 mice lungs was attenuated in those having received the P2712 treatment. Viral replication follows the same tendency. These results directly support the efficacy of the P2712 treatment in preventing viral replication along the course of infection in the Sars-CoV-2/K18-hACE2 model.

Extensive changes in cytokine profiles are documented to be associated with COVID-19 disease progression. The protein levels of 20 different cytokines and chemokines in the lungs, brain and sera were assessed using a multiplex assay (**Figure 6A**). At day 5 post-infection the cytokine response was nearly identical between mice treated with PBS and mock nanoCLAMP P2570, with high levels of pro-inflammatory cytokines and growth factors such as G-CSF, IL-1B, IP-10, M-CSF, MIG, MIP-1a in lungs and brains. The therapeutic nanoCLAMP P2712 had visibly reduced secretion of pro-inflammatory cytokines in the lung and brain, coinciding with the protective effect observed with the clinical score. In plasma samples, this difference is less pronounced and all groups secrete lower levels of pro-inflammatory cytokines. However, compared with serum of PBS treated infected K18-hACE2 animals, a significant reduction of IL6 production after nanoCLAMP P2712 treatment is observed. These data are consistent with cytokine profiling of serum from human patients with COVID-19 and transcriptional analysis of the BAL fluid of human patients, which showed that elevated levels of pro-inflammatory cytokines correlate with disease severity COVID-19 and transcriptional analysis of the BAL fluid of human patients, which showed that elevated levels of pro-inflammatory cytokines correlate with disease severity (Bost et al, 2020, Chen et al, 2020, Yang et al, 2020). Overall, our data suggest that, in the context of the inflammatory response to SARS-CoV-2 in the lungs of K18-hACE2 mice, many cytokines and chemokines are induced, with many having down-regulation patterns of expression after nanoCLAMP P2712 treatment supporting its protective effect in resolution of inflammatory responses.

**Figure 6.**
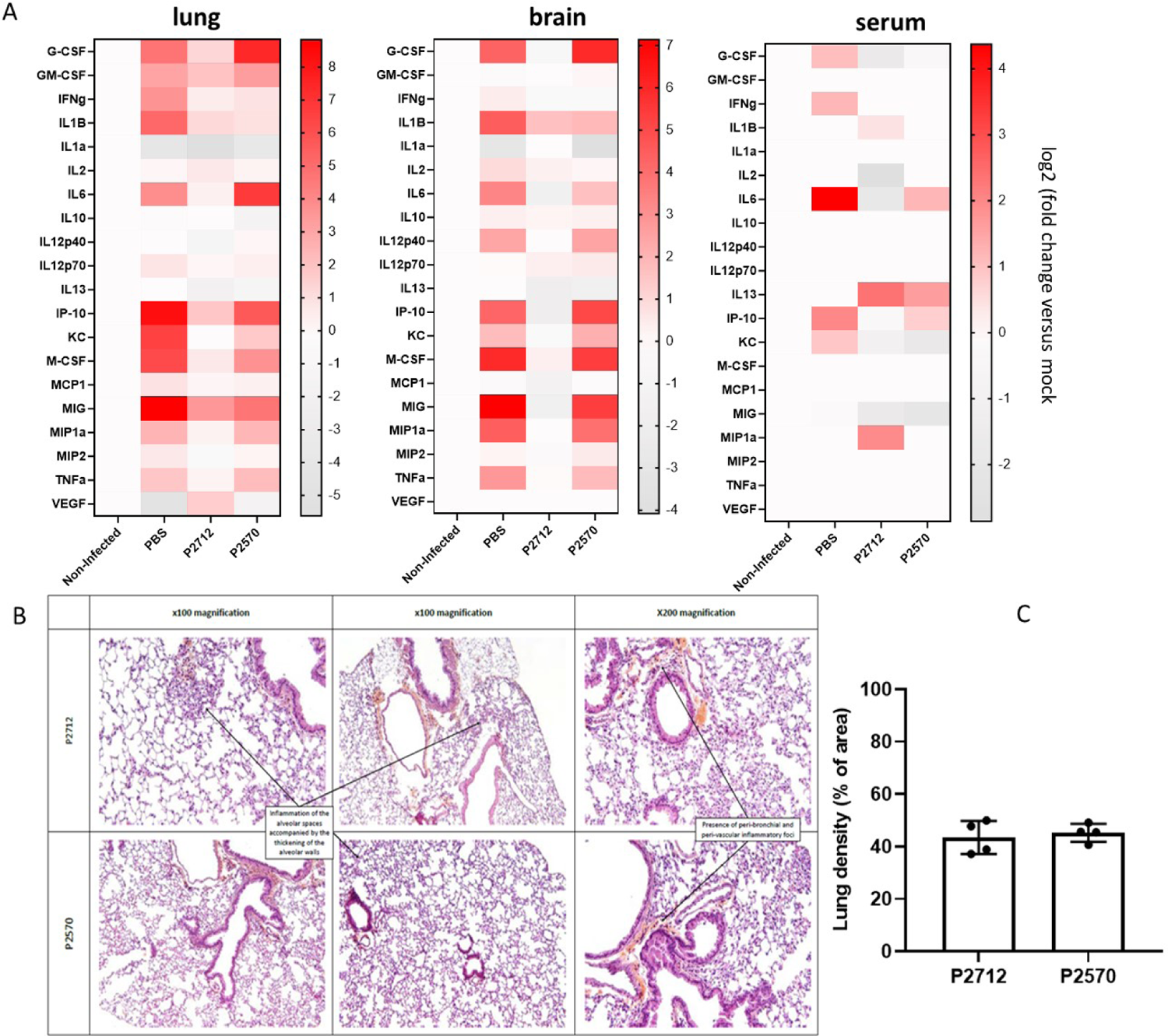
Heat maps of cytokine levels and histopathological analysis of SARS-CoV-2 infection in K18-hACE2 treated miceafter day 5 post infection. (A) (A) Systemic and local cytokine response to SARS-CoV-2 infection in the lung, brain and serum of K18-hACE infected and treated animals by PBS, mock (nanoCLAMP P2570 or therapeutic ligand (nanoCLAMP P2712) at 5-day post infection. For each cytokine, the fold change was calculated as compared with mock-infected animals and the log2[fold change] was plotted in the corresponding heat map. (B) Hematoxylin and eosin (HES) staining of lung sections from infected and treated K18-hACE2 mice following intranasal mock (P2570) or therapeutic nanoCLAMP (P2712) treatment. (C) Lung density analysis on HES stained histological section for representative samples. HES staining (hematoxylikn, eosin, saffron). Nucleus: purple, cytoplasm: pink, muscles: pink-orange, collagen: yellow.

Hematoxylin and Eosin (H&E) staining is the most widely used stain in histology and medical diagnosis. This staining method involves application of hemalum and eosin that color cell nuclei blue and cytoplasm pink to red. The overall patterns of coloration from the stain show the general distribution of cells and provides a histological overview of a tissue structure Analysis of HES lung sections from K18-hACE2 mice infected and treated with nanoCLAMP mock and treatment showed an inflammatory process at 5 dpi (**Figure 6B**). Lung density analysis on HES stained histological section has been done for representative samples. Lung section present peri-bronchial and peri-vascular inflammation accompanied by inflammatory foci in alveolar tissues characterized by local inflammatory cells accumulation and a thickening of the alveolar walls. The measure of lung tissues density in response to SARS-CoV-2 infection showed no decrease or resolution of remodeling between nanoCLAMP P2570 and P2712. This finding corroborates with sustained production of pro-inflammatory cytokines and chemokines at 5 days post infection in the lung, even though down-regulated in compare with K18-hACE2 infected PBS treated animals. In the future, the immune cells infiltrate will need to be analyzed in order to understand better the immune response progression and resolution in the lung upon nanoCLAMP treatment in a dose-response manner.

## 3. Conclusion

Here, effective intranasal therapeutics against SARS-CoV-2 infection in K18-hAce2 mice was achieved using a new class of antibody mimetic, the nanoCLAMP. This work describes the development and initial characterization of a generation of nanoCLAMPs. Initially developed against the Wuhan-Hu-1 RBD, the nanoCLAMP P2712 is a fusion of two nanoCLAMPs that bind simultaneously to the RBD. As variants of concern (VOCs) emerged, the affinity of P2712 as well as its individual subdomains (P2710 and P2609) for the variant RBDs was tracked. It was noted that the affinity of P2609 remained relatively constant for the Wuhan, U.K., S. African, and Delta (7, 6, 10, and 3 nM, respectively) variant, while the P2710 affinity dropped from < 1 pM (Wuhan, UK) to 33 nM (S. African), and 14 nM (Delta) respectively. This resulted in the drop in affinity of P2712 with variants, gradually dropping from < 1 pM (Wuhan, UK) to 7.3 pM (S. African) and 85 pM (Delta) respectively. Since P2710 was able to inhibit Ace2:RBD interaction potently with an IC_50_ = 8 nM, and P2609 only weakly inhibited this interaction (IC_50_ = 573 nM), we hypothesized that mutations affecting P2710:RBD interaction should correlate to lower neutralization rates with pseudovirus variants. Indeed, the pseudovirus neutralization of P2712 for the Wuhan, UK, and S. African variants (IC_50_ = 0.2 to 0.3 nM for U.K and Wuhan) dropped significantly for the S. African variant (IC50 > 100 nM), correlating with the drop-in affinity of subunit P2710 from below 1 pM for Wuhan and U.K RBD to 33 nM for S. African RBD.

With so many variants emerging at the time, we turned our attention to testing actual patient samples to determine whether P2712 could remain effective with increasing RBD mutations. We also wanted to test whether this antibody mimetic could be a candidate for a nasal prophylactic, even if it involved testing using older SARS-CoV-2 strains. The approach here was to build a proof of concept, and if successful, find new neutralizing nanoCLAMPs to more constant regions of the RBD, and repeat.

Since no data exists on the toxicity of nanoCLAMPs, we tested the effect of P2712 on cell viability, and showed that viability gradually declined to about 70% from 0 to 400 µg/ml. Since the IC_50_ of P2712 for pseudovirus was 0.3 nM accounting for 0.01 µg/ml, and viability was near 100% in that range, we concluded P2712 was sufficiently non-toxic to Vero 6 cells to proceed. P2712 indeed neutralized actual patient-derived Wuhan SARS-CoV-2 with an IC_50_ = 0.4 nM using Vero 6 cells, while incubation with P2712 pre- or post-infection did not have any neutralization effect, as expected.

The protection efficacy of P2712 was further validated *in vivo* in K18-hACE2-mice by intranasal administration of P2712 prior to and following infection. By day 9, all of the infected mice not treated with P2712 had succumbed to infection or reached endpoints and were euthanized, whereas 60% of the mice treated with P2712 survived. The viral burden and titer in lung and brain tissues was significantly reduced in P2712 treated vs mock treated animals, demonstrating the protective effect of P2712 on infection of those tissues. Coinciding with these observations was an apparent reduction in several pro-inflammatory cytokines and chemokines in lung and brain, and, to a lesser extent, in serum. In addition, lung tissues inflammation after nanoCLAMP P2570 and 2712 treatment was observed without significant reduction of inflammatory foci and lung density reflecting tissues remodeling. This finding corroborates proinflammatory cytokine and chemokine production that even reduced after nanoCLAMP 2712 treatment is still important to allow progressive resolution of *in situ* lung inflammation detectable by H&E staining. Further analysis of inflammatory lung environment and immune infiltrates need to be performed in order to decipher the mode of action of nanoCLAMP 2712 treatment in control of immune responses upon SARS-CoV-2 infection in lung and brain. The study is currently limited to the use of relatively low dosage of therapeutic, nanoCLAMP (2.25 mg/kg P2712) and compared to an equimolar dose of a single domain, negative control nanoCLAMP P2570. In the future, dose responses will need to be measured to determine the most effective dose of P2712 and its associated clinical, cellular and molecular readout after SARS-CoV-2 2 infection challenge.

SARS-CoV-2 infection of K18-hACE2-transgenic mice supports robust viral replication in the lung, which leads to severe immune cell infiltration, inflammation and pulmonary disease. Thus, the K18-hACE2 mouse is an attractive small animal model for defining the mechanisms of the pathogenesis of severe COVID-19 and may be useful for evaluating countermeasures that reduce virus infection or associated pathological inflammatory responses (Winkler et al, 2020).

Overall, our data suggest that intranasally delivered nanoCLAMP P2712 treatment can protect K18-hACE2 mice against SARS-CoV-2 infection *in vivo*, reduce viral titers in brain and lung and also mitigate the inflammatory response in these tissues. While these preliminary results are encouraging, the potential toxicity and immunogenicity of intranasal administered nanoCLAMP must be explored in more detail in future animal studies. This proof of concept work suggests a path forward for identifying prophylactic and or therapeutic intranasally delivered nanoCLAMP candidates against future SARS-CoV-2 variants by combining individual nanoCLAMPs to more conserved regions of the spike. The rapid generation, facile and low-cost production, and high stability of nanoCLAMPs should make them a useful addition to the growing arsenal of intranasally administered treatments to prevent initial viral host infection.

## 4. Materials and Methods

### 4.1. Identification of nanoCLAMPs specific to SARS-CoV-2 Spike RBD

Nectagen’s nanoCLAMP phage display library NL-21 was panned as described (Suderman et al. 2017) against SARS CoV 2 Spike protein RBD (Sino Biologicals, S1RBD-Fc, cat#40592-VO2H) for rounds 1 and 2, and then biotinylated SARS CoV 2 S protein RBD, His, Avitag (Acro Biosystems, cat# SPD-C82E9), for round 3 (both Wuhan strain). The nanoCLAMPs displayed in this library consist of the nanoCLAMP scaffold with three randomized binding loops, V,W, and Z, which contain 3, 7, and 5 random residues. Loop V residues are comprised of all amino acids except Cys, Glu, Lys, Met, Gln, and Trp. Loops W and Z are comprised of all amino acids except Cys. NL-21 contains over 1.1E10 variants as assessed by the electroporation efficiency of the plasmid library.

For the first round of panning, 2.7 L of 2xYT medium with 2% glucose and 100 mg/ml carbenicillin (2xYT/Glu/CB) was inoculated with 3.6 ml of the NL-21 library glycerol stock (OD_600_ = 75), to an OD_600_ of approximately 0.1, for over 5X coverage of the library size, and grown at 37 °C, 250 rpm until the OD_600_ reached 0.52. The library was infected by adding helper phage VCSM13 (Stratagene, Cat#200251) to 750 ml of culture at an MOI of 20 phage/cell, and incubating at 37°C, 100 rpm for 30 min, then 250 rpm for an additional 30 min. The cells were pelleted at 7500 × g for 10 min, and the media discarded. The cells were resuspended in 1.2 L 2xYT/CB, 70 μg/ml kanamycin (KAN), and incubated 15 h at 30°C, 250 rpm. The cells were combined, and 100 ml was centrifuged at 10k × g for 10 min. The phage containing supernatant was transferred to clean tubes and precipitated by adding 37.5 ml of 5X PEG/NaCl (20% polyethylene glycol 6000/2.5 M NaCl), and incubated on ice for 25 min. The phage was pelleted at 13k × g, 25 min and the supernatant discarded. The phage was resuspended in 10 ml 20 mM NaH_2_PO_4_, 150 mM NaCl, pH 7.4 (PBS), then centrifuged at 15k × g for 15 min to remove insoluble material. The phage was precipitated a second time by adding ¼ volume 5X PEG/NaCl, incubated on ice for 5 min, and pelleted at 13k × g, 10 min at 4°C. The phage pellet was resuspended in 3 ml PBS and quantified by absorbance at 268 nm (A_268_ = 1 for a solution of 5 × 10^12^ phage/ml). Two sets of 100 μl of either Protein A Magnetic Beads (Thermo PI88846) (rounds 1 and 2) or Dynabeads MyOne Streptavidin T1 (ThermoFisher Scientific) (round 3) magnetic beads slurry were washed 2 × 1 ml with PBS-T (PBS with 0.05% Tween 20), applying magnet in between washes to remove the supernatant, and then blocked in 1 ml of 2% dry milk solution in PBS with 0.05% Tween 20 (2% M-PBS-T) for 1 h, rotating, at room temperature. To preclear the phage against beads alone, 1 ml of phage was prepared at a concentration of 2 × 10^13^ phage/ml in 2% M-PBS-T, the block removed from the first set of beads, and the phage added to the beads and incubated 1 h, rotating. The magnet was applied, and the precleared phage removed and transferred to a clean tube. The magnet was applied, and this step repeated two times to ensure no carryover of beads bound to phage to the next step. Fc-RBD (rounds 1,2) was added to the precleared phage to 100 nM final concentration and incubated rotating 1 h. Block was removed from the second set of beads, and the phage/Fc-RBD mix was added to the Protein-A magnetic beads to precipitate the Fc-tagged RBD and bound phage. The beads were washed 8X with PBS-T, 1 ml each, vortexing between each step and applying the magnet. The washed beads were eluted with 800 ul 0.1 M glycine, pH 2.0, 10 min rotating, the magnet applied, and the eluate transferred to 72 ul 2 M Tris base to neutralize. The neutralized phage was then added to 9 ml XL1-blue *E. coli*, which had been grown to OD_600_ = 0.435 and placed on ice. The cells were infected at 37°C, 45 min, 175 rpm, and then expanded to 100 ml 2xYT/Glu/CB and incubated overnight at 30°C, 250 rpm.

The overnight cultures were harvested by measuring the OD_600_, centrifuging the cells at 10k × g for 10 min and then resuspending the cells to an OD_600_ of 75 in 2xYT/18% glycerol. To prepare phage for the next round of panning, 5 ml of 2xYT/Glu/CB was inoculated with 5 μl of the 75 OD_600_ glycerol stock and incubated at 37 °C, 250 rpm until the OD_600_ reached 0.5. The cells were superinfected at 20:1 phage:cell, mixed well, and incubated at 37 °C, 30 min, 150 rpm and then 30 min at 250 rpm. The cells were pelleted at 5500 × g, 10 min, the glucose containing media discarded and the cells resuspended in 10 ml 2xYT/CB/KAN and incubated overnight at 30 °C, 250 rpm. The overnight phage prep was processed as described above. The phage was then prepared at A_268_ = 0.8 in 2% M-PBS-T, and the panning and pre-clearing continued as described, except in the second and third rounds, the target concentration was reduced 10X per round. In round 3, the target was switched to biotinylated RBD and the MyOne T1 streptavidin beads were used. Washes after phage-capture was also increased in the third round, to 12 washes.

### 4.2. Qualitative semELISA of individual clones following panning

At the end of the last panning round, individual colonies were plated on 2xYT/Glu/CB agar plates following the 45 min 150 rpm recovery at 37°C of the infected XL1-blue cells with the eluted phage. The next day, 95 colonies were inoculated into 400 μl 2xYT/Glu/CB in a 96-deep-well culture plate, and grown overnight at 37°C, 300 rpm to generate a master plate, to which glycerol was added to 18% for storage at −80°C. To prepare an induction plate for the ELISA, 5 μl of each master-plate culture was inoculated into 400 μl fresh 2xYT/0.1% glucose/CB medium and incubated for 2.75 h at 37 °C, 300 rpm. IPTG was then added to 0.5 mM and the plates incubated at 30°C with 300 rpm shaking overnight. Because the phagemid contains an amber stop codon, some nanoCLAMP protein is produced without the pIII domain, even though XL1-blue is a suppressor strain, resulting in the periplasmic localization of some nanoCLAMP, of which some percentage is ultimately secreted to the media. The media can then be used directly in an ELISA assay (soluble expression-based monoclonal enzyme-linked immunosorbent assay: semELISA). After the overnight induction, the plates were centrifuged at 1200 × g for 10 min to pellet the cells. Streptavidin coated microtiter plates (ThermoFisher) were rinsed 3 times with 200 μl PBS, and then coated with biotinylated target proteins at 2 μg/ml with 100 μl/well and incubated 1 h. For blank controls, a plate was incubated with 100 μl/well PBS. The coating solution was removed and the plates blocked with 2% M-PBS-T. The block was removed and 50 μl of 4% M-PBS-T added to each well. At this point 50 μl of each induction plate supernatant was transferred to the blank and protein-coated wells and pipetted 10 times to mix, and incubated 1 h. The plates were washed 4 times with 200 μl PBS-T and the plates dumped and slapped on paper towels in between washes. After the washes, 75 μl of 1:2000 dilution anti-FLAG-HRP (Sigma A8592) in 4% M-PBS-T was added to each well and incubated 1 h. The anti-FLAG-HRP was discarded and the plates washed as before. The plates were developed by adding 75 μl TMB Ultra substrate (ThermoFisher), and analyzed for positive signals compared to controls. Positive clones were then grown up from the master plate by inoculating 1 ml 2xYT/2% glucose/100 μg/ml CB with 3 μl glycerol stock and incubated for at least 6 h at 37°C, 250 rpm. The cells were then pelleted and the media discarded. Plasmid DNA was prepared from the pellets using the Qiaprep Spin Miniprep Kit, and the sequences determined by Sanger sequencing at Genewiz (South Plainfield, NJ).

### 4.3. Affinity maturation of nanoCLAMP P2632

A saturation mutagenesis library was constructed using p2632 as template, amplifying the plasmid using mixtures of degenerate primer sets to introduce a single mutation into each variable residue in a loop using the degenerate codon NNK. Degenerate and wildtype primers were mixed such that the library should contain 29% single mutants, 43% double mutants, 22% triple mutants, and 6% wildtype. The amplification of the plasmid and incorporation of variable regions was carried out as described in the library construction scheme in (Suderman et al., submitted). The plasmid library was electroporated into TG1 E.coli and panned for 3 rounds as described. Because the base of this library already had affinity for the target, the concentration of the target protein in round 1 was lowered to 0.1 nM, and was lowered in the two following rounds to 0.01 nM and 0.001 nM, respectively. Also, the wash stringency was increased by increasing the washes from 8 to 12 and including a 7 hour wash in round 2 on the second to last wash. Following the third round of panning, individual clones were assessed by semELISA, and positives characterized as described above for affinity by BLI and monodispersity by SEC.

### 4.4. nanoCLAMP expression and purification

Positive clones identified in the semELISA (above) were subcloned into a pET expression vector containing a N-terminal MGSS-6His tag and either a C-terminal avitag, or no C-terminal tag, depending on the application, and transformed into chemically competent NEc1 *E. coli,* a BL21(DE3) derivative (Nectagen, Inc.). Plasmids were purified using Qiagen miniprep kits (Qiagen) and sequence verified by Sanger sequencing (Genewiz). Glycerol stocks of the plasmids in NEc1 cells were prepared for seeding expression cultures.

Glycerol stocks of NEc1 cells harboring nanoCLAMP expression vectors were used to inoculate 3 ml starter cultures of 2xYT/2% glucose (Glu)/100 mg/ml Carbenicillin (CB) and grown overnight at 37°C, 250 rpm. The overnight cultures were diluted 1:100 into 35 ml of Novagen Overnight Express Instant TB Medium/1% glycerol/CB and incubated 24 h, 30°C, 250 rpm. Cells were pelleted and lysed with 6 M GuHCl, 20 mM Tris, pH 8 (QCB, pH 8), and insoluble material removed by centrifugation at 15k x g, 20 min, 15°C. The cleared lysate was incubated with NiSeph6 FF (Cytiva) for > 1 h rotating, room temperature, then transferred to 2 ml columns. The columns were washed with 6 x 1 ml QCB, pH 8, then refolded with 11 ml of 20 mM MOPS, 150 mM NaCl (MBS), 1 mM CaCl_2_, pH 8. The nanoCLAMPs were eluted with MBS, 1 mM CaCl_2_, 250 mM imidazole, pH 8, buffer exchanged to remove the imidazole using Zeba 7 MWCO desalting columns, and normalized to 1 mg/ml in MBS, 1 mM CaCl_2_, pH 6.5. This procedure was scaled up as necessary to produce larger quantities of protein. For all functional studies, monodisperse monomeric fractions were collected from a Superdex 75 size exclusion column (described below) and pooled.

To generate biotinylated nanoCLAMPs, expression constructs bearing a C-terminal avitag were transformed into chemically competent NEc1/BirA cells (Nectagen, Inc), containing a chloramphenicol selectable plasmid bearing an IPTG-inducible biotin ligase (BirA), and expressed as described above, except the auto-induction culture was supplemented with 50 uM biotin. Prior to eluting the refolded nanoCLAMPs, biotinylation was driven to completion by adding recombinant Mal-BirA (Maltose binding protein with BirA fused to the C-terminus, described in (Li and Sousa 2012)) to 1 uM, ATP to 10 mM, MgCl2 to 10 mM and D-biotin to 150 uM in MBS, pH 8 at a volume of 1.5 ml and incubating rotating at 30C for 2 h. The resin was washed with 5 ml MBS, pH 8, and then 1 ml MBS, pH 8, 1 mM CaCl2 to remove the Mal-BirA. The protein was eluted as described above. Determination of percent biotinylation was measured by incubation of the protein with streptavidin and separating on 12% NuPAGE Bis-Tris (MES buffer) gel and estimating the percentage bound to streptavidin. The protein was buffer exchanged and concentration normalized as described above.

### 4.5. Cloning, expression, and purification of dual epitope nanoCLAMP P2712 and P2712avi

A dual epitope-binding nanoCLAMP (P2712) was constructed by fusing the nanoCLAMPs P2710 to P2609 in the format MGSS-6His-P2710-(GSlinker)-P2609 (see Supporting Information for sequence) or with a C-terminal avitag for biotinylation, in the pET vector described above.

#### P2712 purification from 1 L shake flask for in vivo experiments

The p2712 plasmid was expressed in 1 L autoinduction media as described above, and the pellet lysed with QCB, pH 8 using a polytron. Insoluble material was removed by centrifugation and the cleared supernatant applied to 5 ml NiSeph6 FF, which was washed with 80 ml QCB, pH 8, 25 ml 8 M Urea, 20 mM Tris, pH 8, and then eluted in 8 M Urea, 20 mM Tris, pH 8, 250 mM imidazole. The protein was precipitated with ice cold ethanol, pelleted at 15k x g, 20 min, 4C, and then resuspended with 8 M Urea, 20 mM Tris, pH 8 (Buffer A). The protein was applied to a 1 ml HiTrapQ HP column (Cytiva) and eluted with a gradient of Buffer B (Buffer A with 1 M NaCl) from 0 – 80% B. Fractions containing pure P2712 were pooled and applied to a second IMAC column, washed with 8 M Urea, 20 mM Tris, pH 8, then refolded with MBS, pH 8, 1 mM CaCl2, and finally eluted with refolding buffer with 250 mM imidazole. Monodisperse monomer was collected by size exclusion chromatography on a Superdex 75 column, as described above. Purity was estimated at over 95% by NuPAGE Bis-Tris (MES buffer) and staining with GelCode Blue (Thermo) (SI Appendix, Fig S5).

### 4.6. Endotoxin removal from P2712 and P2570 (neg control)

Endotoxin was removed from purified P2712 and P2570 using the method of Teodorowicz (2017), by extracting with Triton X114 (Sigma) and removing traces of TX114 with BioBeads SM-2 (Bio Rad). Finally, proteins were filtered through a 0.2 um PES filter (Millex-GP SLGPR33RS), and then adjusted to 66.7 uM. Endotoxin levels of each preparation were measured by LAL assay.

### 4.7. Analysis of monodispersity by size exclusion chromatography

Purified nanoCLAMPs were diluted to a final concentration of 0.18 mg/ml in MBS, 1 mM CaCl_2_, pH 6.5, centrifuged at 20k x g, 2 min 4°C, and the supernatants transferred to a clean tube. The samples were loaded into a 125 µl sample loop and injected onto a Superdex 75 10/300 GL column (GE Healthcare Life Sciences, Pittsburg, PA) equilibrated in MBS, 1 mM CaCl_2_, pH 6.5 at a flowrate of 0.65 ml/min. The column was calibrated with Bio-Rad Gel Filtration Standard per manufacturer’s instructions.

### 4.8. Determination of melting temperature by differential scanning fluorimetry

The melting temperature of purified nanoCLAMPs was determined using GloMelt Thermal Shift Protein Stability Kit (Biotium) per manufacturer’s instructions. Briefly, purified nanoCLAMPs were adjusted to 1 mg/ml in MBS, 1 mM CaCl_2_, pH 6.5 and diluted in half with 2X GloMelt (Biotium) and aliquoted to 386 well plate and sealed with optical film. The plate was then heated in a Quantstudio 5 qPCR machine using SYBR Green reporter with no passive reference. The heating profile was 25°C for 2 min; ramp at 0.05°C /sec to 99°C; 99°C for 2 min. T_m_ is defined as the inflection point in the unfolding curve.

### 4.9. Biolayer Interferometry of nanoCLAMPs

Kinetic analysis of interactions between nanoCLAMPs and RBD was carried out on an OctetRed96 using SAX streptavidin coated sensor tips. The tips were transferred first to buffer (MBS, 1 mM CaCl_2_, pH 6.5 + 1% BSA) for 300 sec, then to biotinylated RBD or nanoCLAMP at 2 ug/ml in buffer for 180 sec, then to buffer for 300 sec, then to at least 4 dilutions of nanoCLAMPs or RBDs (depending upon orientation) in buffer (association) for 200 sec, then to buffer (dissociation) for 720 sec. The cells were constantly vortexing at 1000 rpm at rm temp. The kinetics were fit to a 1:1 model and K_d_ calculated using global fit analysis. For high-throughput off-rate analysis, biotinlated RBD was immobilized on SAX tips as described, and then incubated in a single concentration (800 nM) of nanoCLAMP using the parameters above. For epitope binning experiments, SAX sensors were coated with biotinylated P2710avi at 15 uM, then loaded with Wuhan-RBD at 30 nM, and finally incubated with “competitor” non-biotinylated nanoCLAMP at 30 nM, or buffer (neg control).

### 4.10. RBD/Ace2 competitive inhibition ELISA

Polystyrene Protein A coated microtiter plates (ThermoFisher: 15132) were coated by incubation on a plate shaker at room temperature with 0.5 µg/mL of recombinant SARS-CoV-2 Spike RBD-Fc (SinoBiological: 40592-V02H). Negative control wells were incubated with 2% Milk + PBS-T (20 mM NaH2PO4, 150 mM NaCl (PBS) + 0.05% Tween 20) or 0.5 µg/mL recombinant SARS-CoV-2 Spike RBD-Fc. Plates were washed 3 times with PBS-T. Following 1 h incubation with recombinant SARS-CoV-2 Spike RBD-Fc, preparations of 800 nM nanoCLAMP candidates were serially diluted 4-fold (i.e. 800, 200, 50, 12.5, 3.125, 0.781, and 0.195 nM) with Biotinylated Human ACE2/ACEH Protein, His, Avitag (AcroBiosystems: AC2-H82E6) containing dilution buffer at a final Biotinylated Human ACE2 concentration of 25 ng/mL. The solutions were incubated on a plate shaker at room temperature for 1 h. Biotinylated Human ACE2 containing dilution buffer was added to the negative control wells. Following the 1 h incubation, the wells were washed 3 times with PBS-T. All wells were then incubated Streptavadin-HRP (ThermoSci: N504) at a 1:5000 dilution on a plate shaker at room temperature for 1 h. After incubation with SA-HRP, the plate was washed 3 times with PBS-T and developed with 1-Step Ultra TMB-ELISA Substrate Solution (FisherSci: 34028) for approximately 7.5 min. The reaction was stopped using 2N Sulfuric Acid solution. Absorbance was read using Molecular Devices, Kinetic Microplate Reader and analyzed using SoftmaxPro 5.4.

### 4.11. Pseudovirus neutralization assay

Lentivirus pseudo-typed with Spike protein from SARS-CoV-2 Wuhan-Hu-1 and bearing a Luciferase-IRES-ZsGreen payload, was prepared as described (Crawford et al 2020) and also purchased from Integral Molecular (Cat# RVP-701L). Lentivirus pseudotyped with UK Variant (B.1.1.7) and South African Variant (B.1.351) Spike proteins were purchased from Integral Molecular (cat# RVP-706L and RVP-707L, respectively). Ace2-expressing HEK293T cells (293T-hsACE2) were purchased from Integral Molecular (Cat# C-HA102). Briefly, 293T-hsAce2 cells were plated in 96-well format at 2.0 × 104 cells/well. On the day of infection, the pseudo-typed viruses bearing luciferase reporter were pre-incubated with the serially diluted nanoCLAMPs in DMEM containing 10% FBS, 100 units/mL of penicillin and streptomycin at 37°C for 1 h and applied to the cells. Luminescence was measured 72 h post-infection using the Bright-GloTM Assay System (Promega: E2610) when assaying in-house generated Wuhan pseudovirus, or Renilla-GloTM Assay System (Promega: E2710) when assaying the Integral Molecular pseudoviruses. Luminescence was read using Molecular Devices, Kinetic Microplate Reader with 1000 ms integration time and analyzed using SoftmaxPro 5.4. Psuedo-typed viruses were diluted to yield a Signal to Background ratio of approximately 500-fold.

### 4.12. Fluorescent labelling of nanoCLAMP P2717LC

nanoCLAMP-LC (1 µM) was mixed with Tris(2-carboxyethyl)phosphine (Sigma, France) at 10 µM in 100 µL for 20 min at room temperature. Then 10 µL of CF-488A-Maleimide (Sigma, France) was added to the mixture and incubated over night at room temperature. The formed NanoCLAMP-Dye was purified with Zeba™ Spin Desalting Columns (Thermofisher, France) 4 times, following supplier procedures. The success of the fluorescence lableinw was evaluated with Nanodrop.

### 4.13 In vitro experiments

#### Vero E6 cells

(ATCC CRL-1586) were cultured in Dulbecco’s modified Eagle medium (DMEM) supplemented with 10% fetal bovine serum (FBS), 1% L-glutamine, 1% antibiotics (100 U mL^−1^ penicillin, and 100 μg mL^−1^ streptomycin), in a humidified atmosphere of 5% CO_2_ at 37 °C.

#### Cell Viability

Vero E6 Cells were seeded in 96-well plates (2 × 10^4^ cells/well) 24 h before test. The culture medium was replaced with a fresh medium that contains increasing concentrations of NanoCLAMP for 72 h from 0 to 400 μg mL^−1^. The old medium was then aspirated, and cells were washed with PBS. The cell viability was evaluated using resazurin cell viability assay. Briefly, 100 mL of the resazurin solution (11 μg mL^−1^) in DMEM/10% FBS was added to each well, and the plate was incubated for 4 h. The fluorescence emission of each well was measured at 593 nm (20 nm bandwidth) with an excitation at 554 nm (18 nm bandwidth) using a Cytation 5 Cell Imaging Multi-Mode Reader (BioTek Instruments SAS, France). Each condition was replicated three times, and the mean fluorescence value of cells treated the same way in the absence of nanoCLMAPs was taken as 100% cellular viability

#### Flow cytometry

Vero E6 Cells were seeded in 6-well plates (2 × 10^6^ cells/well) 24 h before the test. The culture medium was replaced with a fresh medium with 25.5 nM of NanoCLAMP-Dye for 4 hours and cells were washed with PBS1x. The fluorescence emission of each well was measured at 530 nm (30 nm bandwidth) with excitation at 488 nm (18 nm bandwidth) using an Attune NxT (Invitrogen, France) for each condition we measure 50.000 events.

#### Virus Amplification and Titration

Vero E6 cells were plated in 96-well plates (5 × 10^5^ cells/well) 24 h before performing the virus titration. A clinical isolate, obtained from a SARS-CoV-2 positive specimen, was cultured on Vero E6 cells with MOI of 145 (infectious agents to infection targets: 7.27×10^7^ / 5.00 ×10^7^). Infected cell culture supernatant was centrifuged for 5 min at 5 000 rpm at 4°C to obtain a virus suspension. Virus suspensions were distributed in 6 wells in DMEM supplemented with 10% FBS (Fetal Bovine Serum) to Vero E6 cells, 1% antibiotics (100 U/mL penicillin, and 100 μg/mL streptomycin), and 1% L-glutamine. The plates were incubated for 5 days in 5% CO_2_ atmosphere at 37 °C.

#### SARS-CoV-2 RNA extraction

Viral RNA was extracted from supernatant of three replicate of each samples with the QIAmp Viral RNA kit (QIAGEN, USA).

#### SARS-CoV-2 RT-PCR

The Luna Universal Probe One-Step RT-qPCR kit (New England Biolabs, USA) was used. The primers used are M-475-F and M-574-R (Eurofins Genomics, Germany) (Toptan et al, 2020). The thermal cycling conditions were as follows: 10 min at 55 ◦C for lysis and reverse transcription, 1 min at 95 °C for polymerase activation and 45 cycles of 10 s at 95 ^°^C, 30 s at 60 ^°^C (data acquisition) and a melt curve acquisition with 1 min at 95°C, 30 s at 60°C and 30 sec at 95 °C. Undetectable SARS-CoV-2 levels were set to Ct 40. Amplification was performed on Mx3000P QPCR System (Agilent Technologies, USA). (Kriegova et al, 2020)

### 4.14. Mechanistic investigations

Vero E6 cells were seeded at a density of 5 × 10^5 cells per well in 96-well plates and allowed to attach for 24 hours before testing.

i. The culture medium (DMEM supplemented with 10% FBS, 1% L-glutamine, and 1% antibiotics consisting of 100 U/mL penicillin and 100 μg/mL streptomycin) was replaced with fresh medium containing 60 nM of NanoClamp for 30 minutes. The cell layers were then washed 5 times with 1X PBS and the viral solution was incubated with the cells for 30 minutes. The cell layers were washed another 5 times with 1X PBS and the plates were incubated for 6 days at 37°C with 5% CO2.
ii. A solution of NanoClamp and virus was prepared in culture medium and incubated for 30 minutes. The cell layers were then incubated with this solution for 30 minutes. After incubation, the cell layers were washed 5 times with 1X PBS and the plates were incubated for 6 days at 37°C with 5% CO2.
iii. A viral solution was diluted 10-fold with culture medium and incubated with the cell layer for 30 minutes. The cell layers were then washed 5 times with 1X PBS and incubated with fresh medium containing 60 nM of NanoClamp for an additional 30 minutes. The cell layers were washed another 5 times with 1X PBS and the plates were incubated for 6 days at 37°C with 5% CO2.

In all three cases, the plates were monitored daily using an inverted microscope (ZEISS Primovert) to assess the extent of virus-induced cytopathic effects in the cell culture.

### 4.15. SARS-CoV-2 infection and treatment in mice

Animal housing and experimental procedures were conducted according to the French and European Regulations (Parlement Européen et du Conseil du 22 Septembre 2010, Decret n° 2013-118 du 1er fevrier 2013 relatif a la protection des animaux utilisées a des fins scientifiques) and the National Research Council Guide for the Care and Use of Laboratory Animals (National Research Council (U. S.), Institute for Laboratory Animal Research (U.S.), and National Academies Press (U.S.), Eds., Guide for the care and use of laboratory animals, 8th ed. Washington, D.C: National Academies Press, 2011). The animal BSL3 facility is authorized by the French authorities (Agreement N° B 13 014 07). All animal procedures (including surgery, anesthesia, and euthanasia, as applicable) used in the current study were submitted to the Institutional Animal Care and Use Committee of the CIPHE and approved by the French authorities (APAFIS#26484-2020062213431976 v6). All CIPHE BSL3 facility operations are overseen by a biosecurity/biosafety officer and accredited by the Agence Nationale de Sécuritée du Médicament (ANSM).

#### Animals

Heterozygous K18-hACE C57BL/6J mice (strain: 2B6. Cg-Tg (K18-ACE2)2Prlmn/J) were obtained from The Jackson Laboratory. All breeding, genotyping, and production of K18-hACE2 was performed at the CIPHE. The sample size was based on previous articles reporting the use of K18-hACE2 mice in SARS-CoV2 challenge experiments (10 animals per experimental group). Animals were housed in groups within cages and fed with standard chow diets.

#### Wuhan/D614 SARS-CoV-2 virus production

Vero E6 cells (CRL-1586; American Type Culture Collection) were cultured at 37 °C in Dulbecco’s modified Eagle’s medium (DMEM) supplemented with 10% fetal bovine serum (FBS), 10 mM HEPES (pH 7.3), 1 mM sodium pyruvate, 1X non-essential amino acids, and 100 U/ mL penicillin/streptomycin. The strain Beta CoV/France/IDF0372/2020 was supplied by the National Reference Centre for Respiratory Viruses hosted by the Institut Pasteur (Paris, France). The human sample from which strain BetaCoV/France/ IDF0372/2020 was isolated was provided by the Bichat Hospital, Paris, France. Infectious stocks were grown by inoculating Vero E6 cells and collecting supernatants upon observation of the cytopathic effect. Debris was removed by centrifugation and passage through a 0.22 mm filter. Supernatants were stored at 80 °C.

#### Infection assay of K18-hACE2 transgenic mice

Intranasal virus and anti-viral treatment were performed under anesthesia and all efforts were made to minimize animal suffering. Eight to twelf week old heterozygous K18-hACE C57BL/6J mice (strain: 2B6. Cg-Tg (K18-ACE2)2Prlmn/J) were inoculate with 2.5×10^4^ PFU Wuhan/D614 SARS-CoV-2 via intranasal administration of 30 µL. Daily treatments were administered intranasally at 5 mg kg^-1^ of 2.5 mg kg^-1^ using the average weight of each groups (30 µl volume). Mice were monitored daily for morbidity (body weight) and mortality (survival). During the monitoring period, mice were scored for clinical symptoms (weight loss, eye closure, appearance of the fur, posture, and respiration). Mice obtaining a clinical score defined as reaching the experimental end-point were humanely euthanized.

#### Measurement of SARS-CoV-2 viral load by RT-qPCR and TCID50 (50% of tissue-culture infective dose)

For viral titration by RT-qPCR, tissues were homogenized with ceramic beads in a tissue homogenizer (Precellys, Bertin Instruments) in 0.5 mL RLT buffer. RNA was extracted using the RNeasy Mini Kit (QIAGEN) and reverse transcribed using the High-Capacity cDNA Reverse Transcription Kit (Thermo Fisher Scientific). Amplification was carried out using OneGreen Fast qPCR Premix (OZYME) according to the manufacturer’s recommendations. The number of copies of the SARS-CoV-2 RNA-dependent RNA polymerase (RdRp) gene in samples was determined using the following primers: forward primer catgtgtggcggttcactat, reverse primer gttgtggcatctcctgatga. This region was included in a cDNA standard to allow the copy number determination down to »100 copies per reaction. The copies of SARS-CoV-2 were compared and quantified using a standard curve and normalized to total RNA levels. An external control (mock-infected wildtype animal, nondetectable in the assay) and a positive control (SARS-CoV-2 cDNA containing the targeted region of the RdRp gene at a concentration of 10^4^ copies/µl [1.94 £ 10^4^ copies/µl detected in the assay]) were used in the RT-qPCR analysis to validate the assay. The median tissue-culture infectious dose (TCID50) represents the dilution of a virus-containing sample at which half of the inoculated cells show signs of infection. To perform the assay, lung and brain tissues were weighed and homogenized using ceramic beads in a tissue homogenizer (Precellys Bertin Instruments) in 0.5 ml RPMI media supplemented with 2% FCS and 25 mM HEPES. Tissue homogenates were then clarified by centrifugation and stored at 80 °C until use. Forty-thousand cells per well were seeded in 96-well plates containing 200 µl DMEM +4% FCS and incubated for 24 h at 37 °C. Tissue homogenates were serially diluted (1:10) in RPMI media and 50 µl of each dilution was transferred to the plate in six replicates for titration at five-days post-inoculation. Plates were read for the CPE (cytopathology effect) using microscopy reader and the data were recorded. Viral titers were then calculated using the Spearman-Karber formula and expressed as TCID50/mg of tissue.

#### Histology analysis

At day 5 post infection, 3 mice per group were euthanatized, lung and brain tissues harvested and fixed in Sterilin tubes with 4% paraformaldéhyde. Fixed left lung (4% paraformaldehyde) were paraffin-embedded (Leica Pearl®). Three Three-µm transversal sections were cut, mounted on Superfrost glass slides (Fischer Scientific), and stained with hematoxylin eosin saffron (HES). To determine the severity of tissue remodeling observed on histological sections, area of stained tissue was measured and reported to the total section area. Measure were realized with the Cell Dimension® software coupled to a DP72 camera (Olumpus) on three sections for each animal.

#### Cytokine and chemokine protein measurements

K18-hACE2 transgenic mice were infected (intranasally challenged with 2.5 x 10^4^ PFU of SARS-CoV-2 Wuhan) and treated according to administration regimen by intranasal inoculation of P2712 or P2570 (6 hours before infection and day 1 and 2 post-infection). At day 5 post infection, blood was collected from a submandbulary sampling and plasma were isolated after centrifugation. Lung and brain were harvested in Precellys® tubes containing RPMI medium completed with Pen/Strep, Hepes and FCS than homogeneized with the Precellys®. Then the samples were inactivated by a mix of Triton 10X and RPMI medium at a final concentration of 0.5% Triton. A kit, created by Merck Millipore, in order to cover the overall cytokine and chemokine responses was used here. This kit coupled with the Luminex® platform in a magnetic bead format provide the advantage of ideal speed and sensitivity, allowing quantitative multiplex detection of dozens of analytes simultaneously, which can dramatically improve productivity. It contained cytokines standards to make standard curve and to ensure lot-to-lot consistency, two quality control to qualify assay performance, detection antibody cocktails designed to yield consistent analyte profiles within panel, streptavidin-PE and a panel of magnetic beads that recognize each one of the following analytes; IL-1a, IL-1b, IL-2, IL-6, IL-10, IL12p40, IL-12p70, IL-13, M-CSF, G-CSF, GM-CSF, IP-10 (CXCL10), IFN-g, MCP-1 (CCL2), MIP 1-a (CCL3), TNF-a, VEGF, CXCL9 (MIG), CXCL1 (KC), MIP-2. In a 96-wells plate previously washed with wash buffer, we incubated overnight (at 4°C and on shaker) and according a template, the standards, the quality controls and the samples (plasma and organs supernatants) with assay buffer and magnetic beads panel. The following day the plate was washed twice with wash buffer, then we added the detection antibodies cocktail and incubate 1h at room temperature on shaker before we added the Streptavidine-PE and incubate for 30min. Then the plate was washed twice and the samples were suspended in assay buffer for 5 min on shaker before reading on MagPix® Instrument.

### 4.16. Statistics

Graphpad Prism software version 8 was used for non-parametric statistics and plots, as described in the figure legends. Heatmaps were generated using the heatmap function from package NMF in R software, version 4.0.0. R: A language and environment for statistical computing. R Foundation for Statistical Computing, Vienna, Austria. URL: https://www.R-project.org. Statistical differences in the expression of standardized bio-markers were determined using the nonparametric Wilcoxon test, adjusting for multiple testing using the Benjamini & Hochberg correction.

## Data Availability

all data are available opn request

## Acknowledgments

Financial support from the Centre National de la Recherche Scientifique (CNRS) and the University of Lille as well as ANR FLU is acknowledged. We also thank Cathleen Lutz and The Jackson Laboratory for providing the K18-hACE2 mice and Pr. Sylvie van der Werf, Dr. X.Lescure, and Pr. Y. Yazdanpanah for the BetaCoV/France/IDF0372/2020 strain and François Daubeuf UAR3286 – PCBIS Plate-forme de Chimie Biologique Intégrative de Strasbourg Technologie du médicament (Techmed’ILL) for histology analysis.

## References

Bost P, Giladi A, Liu Y, Bendjelal Y, Xu G, et al. (2020) Host–viral infection maps reveal signatures of severe COVID-19 patients. Cell 181: 1475–1488.

Cameroni E, Bowen JE, Rosen LE, Saliba C, Zepeda SK, et al. (2022) Brodaly neutralizing antibodies overcome SARS-CoV-2 omicron antigetnic shift. Nature 602: 664.

Case JB, Che n, R. E., Cao L, Ying B, Winkler ES, et al. (2021) Ultrapotent miniproteins targeting the SARS-CoV-2 receptor-binding domain protect against infection and disease. Cell Host & Microbe, 2 29: 1–11.

Chen X, Zhao B, Qu Y, Chen Y, Xiong J, et al. (2020) Detectable serum SARS-CoV-2 viral load (RNAaemia) is closely correlated with drastically elevated interleukin 6 (IL-6) level in critically ill COVID-19 patients. Clin. Infect. Dis. 71: 1937–1942.

Chi X, Yan R, Zhang J, Zhang Y, Hao M, et al. (2020) A neutralizing human antibody binds to the N-terminak domain of the spike protein of SARS-CoV-2. Science 369: 650–655.

Chonira V, Young D, Kwon YD, Jason Gorman J, James Brett Case JB, Zhiqiang Ku Z, et al. (2023) A potent and broad neutralization of SARS-CoV-2 variants of concern by DARPins. Nat. Chem. Biol. 19: 284-291.

Dai L, Gao GF. (2020) Viral targets for vaccines against COVID-19. Nat. Rev. Immunol. 21: 73–82

Güttler T, Aksu M, Dickmanns A, Stegmann KM, Gregor K, et al. (2021) Neutralization of SARS-CoV-2 by highly potent, hyperthermostable, and mutation-tolerant nanobodies. EMBO journal 2021 40: e107985.

Hadjichrysanthou C, Beukenhorst AL, Koch CM, Alter G, Goudsmit J, et al. (2022) Exploring the Role of Antiviral Nasal Sprays in the Control of Emerging Respiratory Infections in the Community. Infect. Dis. Ther. 11: 2287–2296.

Hoffmann M, Kleine-Weber H, Schroeder S. (2020) SARS-CoV-2 cell entry depends on ACE2 and TMPRSS2 and is blocked by a clinically proven protease inhibitor. CEll 271– 280: e8.

Imai M, Mutsumi Ito M, Kiso M, Yamayoshi S, Uraki R, et al. (2023) Efficacy of Antiviral Agents against Omicron Subvariants BQ.1.1 and XBB. New. Engl. J. Med. 388: 89-91.

Kriegova E, Fillerova R, Kvapil P. (2020) Direct-RT-qPCR Detection of SARS-CoV-2 without RNA Extraction as Part of a COVID-19 Testing Strategy: From Sample to Result in One Hour. Diabgostics 10: 605.

Ku Z, Xie X, Hinton PR, Liu X, Ye X, et al. (2021) Nasal delivery of an IgM offers broad protection from SARS-CoV-2 variants. Nature 595: 718–723.

Kuramochi T, Gan SW, Ho AWS, Wang B, Kageji N, et al. (2022) Comprehensive enginnering of a therapeytic neutralizing antibody targeting SARS-CoV-2 spike protein to neutralie escepae variantes. MABS 14: e2040350.

Li H, Zhong D, Luo H, Shi W, Xie S, et al. (2022) Nanobody-based CAR T cells targeting intracellular tumor antigens. *Front*. Immunology. 156: 113919.

Lin Y, Shuai Yue S, Yang Y, Yang S, Pan Z, et al. (2023) Nasal Spray of Neutralizing Monoclonal Antibody 35B5 Confers Potential Prophylaxis Against Severe Acute Respiratory Syndrome Coronavirus 2 Variants of Concern: A Small-Scale Clinical Trial. Clin. Infect. Dis. 76: e336–e341.

Puray-Chavez M, LaPak KM, Schrank TP, Elliott JL, Bhatt DP, et al. (2021) Systematic analysis of SARS-CoV-2 infection of an ACE2-negative human airway cell Cell Rep. 36: 109364.

Pymm P, Adair A, Chan J-L, Cpnney JP, Mordant FL, et al. (2021) Nanobody cocktails potently neutralize SARS-CoV-2 D614G N501Y variant and protect mice. Proc. Natl. Acad. Sci. U.S.A 118: e2101918118.

Ratcliffe NA, Castro HC, Gonzalez MS, Mello CB, Dyson P. (2022) Reaching the Final Endgame for Constant Waves of COVID-19. Viruses 14: 2637.

Ratcliffe NA, Castro HC, Paixão IC, Evangelho VGO, Azambuja P, et al. (2021) Nasal therapy—The missing link in optimising strategies to improve prevention and treatment of COVID-19. PLoS Pathogens 17: e1010079.

Renn A, YFu Y, Hu X, Hall MD, Simenonov A. (2020) Fruitful Neutralizing Antibody Pipeline Brings Hope To Defeat SARS-Cov-2. Trends Pharmacol. Sci. 41: 815–829.

Sécher T, Bodier-Montagutelli E, Parent C, Bouvart L, Cortes M, et al. (2022) ggregates Associated with Instability of Antibodies during Aerosolization Induce Adverse Immunological Effects. Pharmaceutics 14: 671.

Suderman R, Rice DA, Gibson SD, Strick EJ, Chao D. (2017) Development of polyol-responsive antibody mimetics for single-step protein purification. Protein Expr. Purif. 134: 114–124.

Titong A, Kankanamalage SG, Dong J, Huang B, Spadoni N, et al. (2022) First-in-class trispecific VHH-Fc based antibody with potent prophylactic and therapeutic efficacy against SARS-CoV-2 and variants. Sci. Rep. 12: 4163.

Toptan T, Hoehl S, Westhaus S, Bojkova D, Berger A, et al. (2020) Optimized qRT-PCR Approach for the Detection of Intra- and Extra-Cellular SARS-CoV-2 RNAs. Int. J. Mol. Sci. 21: 4396.

Wang Q, Iketani S, Li Z, Liu L, Guo Y, et al. (2023) Alarming antibody evasion properties of rising SARS-CoV-2 BQ and XBB subvariants. CEll 279–28: e8.

Weinrich DM, Sivapalasingam S, Norton T, Ali S, Gao H, et al. (2020) REGN-CoV2, as neutralizing antibody cocktail, in outpatients with Covid-19. N. Eng. J. Med. 384: 283–251.

Winkler ES, Bailey AL, Kafai NM, Nair S, McCune BT, et al. (2020) SARS-CoV-2 infection of human ACE2-transgenic mice causes severe lung inflammation and impaired function. Nat. Immunol. 21: 1327–1335.

Wu X, Cheng L, Fu M, Huang B, Zhu L, et al. (2021) A potent bispecific nanobody protects hACE2 mice against SARS-CoV-2 infection via intranasal administration. 37: 109869.

Wu X, Wang Y, Cheng L, Ni F, Zhu L, et al. (2022) Short-Term Instantaneous Prophylaxis and Efficient Treatment Against SARS-CoV-2 in hACE2 Mice Conferred by an Intranasal Nanobody (Nb22). Front. Immunol 13: 865401.

Yang Y, Shen C, Li J, Yuan J, Wei J, et al. (2020) Plasma IP-10 and MCP-3 levels are highly associated with disease severity and predict the progression of COVID-19. J. Allergy Clin. Immunol. 146: 119–127.

